# Improvement in auditory spatial discrimination from ambiguous visual stimuli is not explained by ideal observer causal inference

**DOI:** 10.1101/598425

**Authors:** Madeline S. Cappelloni, Sabyasachi Shivkumar, Ralf M. Haefner, Ross K. Maddox

**Affiliations:** Biomedical Engineering, University of Rochester, Rochester, NY; Del Monte Institute for Neuroscience, University of Rochester, Rochester, NY; Brain and Cognitive Sciences, University of Rochester, Rochester, NY; Center for Visual Science, University of Rochester, Rochester, NY; Neuroscience, University of Rochester, Rochester, NY

## Abstract

In order to survive and function in the world, we must understand the content of our environment. This requires us to gather and parse complex, sometimes conflicting, information. Yet, the brain is capable of translating sensory stimuli from disparate modalities into a cohesive and accurate percept with little conscious effort. Previous studies of multisensory integration have suggested that the brain’s integration of cues is well-approximated by an ideal observer implementing Bayesian causal inference. However, behavioral data from tasks that include only one stimulus in each modality fail to capture what is in nature a complex process. Here we employed an auditory spatial discrimination task in which listeners were asked to determine on which side they heard one of two concurrently presented sounds. We compared two visual conditions in which task-uninformative shapes were presented in the center of the screen, or spatially aligned with the auditory stimuli. We found that performance on the auditory task improved when the visual stimuli were spatially aligned with the auditory stimuli—even though the shapes provided no information about which side the auditory target was on. We also demonstrate that a model of a Bayesian ideal observer performing causal inference cannot explain this improvement, demonstrating that humans deviate systematically from the ideal observer model.

## Introduction

As we navigate the world, we gather sensory information about our surroundings from multiple sensory modalities. Information gathered from a single modality may be ambiguous or otherwise limited, but by integrating information across modalities, we form a better estimate of what is happening around us. While our integration of multisensory information seems effortless, the challenge to the brain is non-trivial. The brain must attempt to determine whether incoming information originates from the same source, as well as estimate the reliability of each modality’s cues so that they may be appropriately weighted.

Studies of multisensory integration have explained how a Bayesian ideal observer could solve this problem by combining reliability-weighted evidence from multiple sensory modalities. In the forced integration model, an observer gathers evidence from multiple modalities and combines them according to the modality’s reliability [1]. Importantly this allows for the most reliable sensory estimate to dominate the percept while noisier measurements have less influence; however, it also implies that percepts of distinct stimuli that in actuality originate from independent sources must nonetheless be perceptually influenced by each other. More recently, causal inference has expanded upon the forced integration model by allowing the observer to treat stimuli as originating from different sources. The observer first determines whether both pieces of evidence are likely to come from a common source, and if so weight them by their reliabilities as in the forced integration model, to generate a combined percept [2]. In their basic forms neither model attempts to contend with scenes more complex than a single stimulus in each modality.

Numerous experiments have shown that humans behave as ideal or near-ideal Bayesian observers performing forced integration [3–6] or causal inference [7–10]. There have even been efforts to reveal which brain structures contribute to Bayesian computations [11, 12]. However, those studies have not considered scenarios in which many sources in an environment give rise to multiple cues within each modality. Here we test the Bayesian causal inference model using a new paradigm, and in doing so introduce a key question missing from these prior studies, but common in the natural world; which auditory and visual stimuli will be integrated when multiple stimuli exist in each modality?

In the case of a single stimulus in each modality, visual influence on auditory location has been largely demonstrated by studies of perceptual illusions. Notably the ventriloquist effect, a bias of auditory location toward visual location when cues of both modalities are presented simultaneously [13], has been extensively characterized. The influence of the visual location depends mainly on two factors: the discrepancy between the two stimuli (with visual-induced bias waning as the spatial separation becomes too large) [14], and the size of the visual stimulus (smaller, more reliable, visual stimuli yielding a larger bias) [4]. Dependence on discrepancy points to a causal inference structure, while size dependence indicates a weighting by the quality of the location estimates (larger visual stimuli are localized less accurately). Agreement with the Bayesian causal inference model [2, 4] would indicate that the bias is due to an integration of the two cues in which the brain produces a combined estimate of location. Therefore, congruent auditory and visual evidence should result in a more accurate estimate of object location than auditory evidence alone.

In this study we engaged listeners in a concurrent auditory spatial discrimination task to look for a benefit from spatially aligned, task-uninformative visual stimuli. We presented two sounds, a tone and noise, with centrally located or spatially aligned visual stimuli of per-trial random color and shape. Listeners were asked to report which side the tone was on. Importantly, those shapes do not provide information about the correct choice in either condition, only providing information about the separation of the two auditory stimuli in the spatially aligned condition. We investigated whether subjects nonetheless benefited from this additional information and improve their performance on the task as one might predict from an extrapolation of the ventriloquist effect. Our results show a benefit due to the spatially aligned shapes. However, an extension of the ideal Bayesian causal inference model for two auditory and two visual stimuli could not explain any difference in auditory performance between the two visual conditions.

## Results

### Psychophysics

We engaged listeners in an auditory spatial discrimination task to see if they could benefit from spatially aligned task-uninformative visual stimuli. Listeners were presented with two simultaneous sounds (a tone complex and noise token with the same spectral shape) localized symmetrically about zero degrees azimuth and asked to report which side the tone was on. Concurrently, two *task-uninformative* visual stimuli of per-trial random shape and hue were presented. In two conditions (Figure 1), visual stimuli were either spatially aligned with the auditory stimuli (“Matched” condition) or in the center of the screen (“Central” condition) as a control. For both conditions, auditory separations ranged from 1.25 degrees to 20 degrees. We measured the improvement in performance due to the spatially aligned shapes as the difference in percent correct between matched and central conditions for each separation (Figure 2A). Averaging across separations, the improvement was significant with (*p* = 0.007, *t*-test). The effect was individually significant at moderate and large separations (5 degrees (*p* = 0.003) and 20 degrees (*p* = 0.02)).

**Fig 1.**
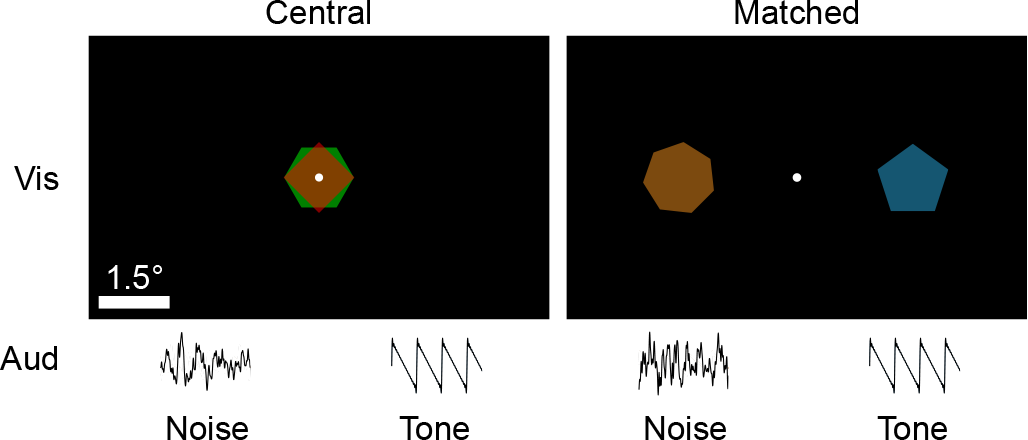
Listeners fixate while concurrently hearing two auditory stimuli on either side of the fixation dot and seeing two random shapes that are either centrally located or aligned spatially with the auditory stimuli. Shapes are presented in alternating frames to avoid overlap.

**Fig 2.**
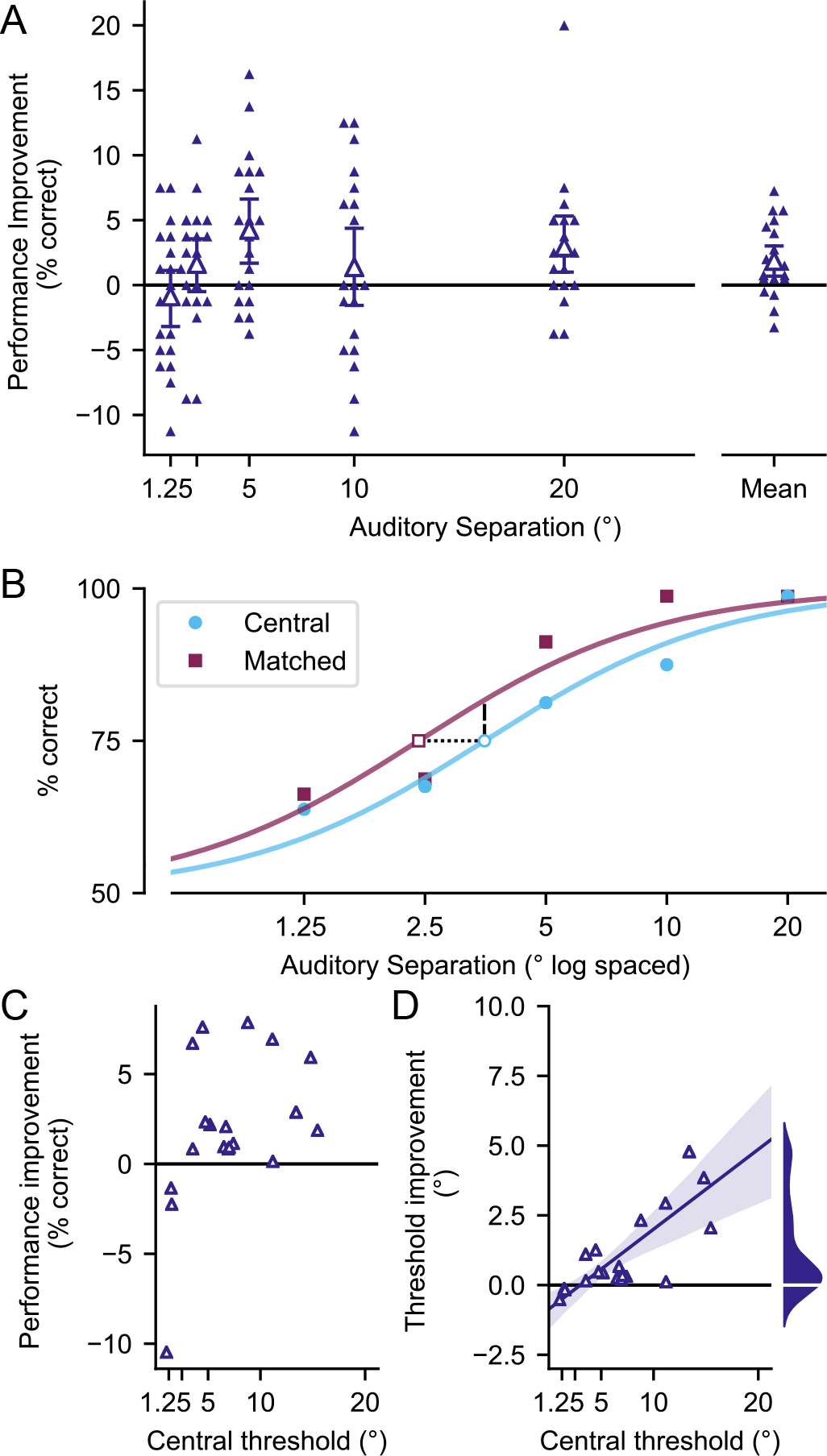
Behavioral results comparing central and matched conditions. **A)** Improvement in performance at each angle averaged across subjects. Error bars show 95% confidence intervals and individual subjects shown as small triangles. **B)**. Sigmoidal fits of the data in log units for a single subject who shows the effect. **C)**. Improvement in performance at each subject’s separation threshold in the central condition (dashed line in B). **D)** Improvement in threshold for each subject (dotted line in B). Line of best fit and 95% confidence intervals also shown. Marginal distribution of threshold improvement shown to the right. There is more mass towards positive threshold improvement than negative.

To further understand the effect, we calculated 75% thresholds for each condition by fitting a psychometric function to each subject’s response data (Figure 2B). Improvements in threshold across conditions and improvements in performance at threshold are shown in Figure 2C-D. A decrease in separation thresholds (dashed line Figure 2B) is necessarily paired with an increase in percent correct at threshold (dotted line Figure 2B) due to the fit method (slope and lapse rate of the sigmoid were determined from responses to both conditions and only the threshold parameter of the function was allowed to differ between the two conditions). Nonetheless, we find that improvements at threshold (and consequently, performance at threshold) are significant across the population (*p* = 0.0002, sign test). The average threshold improvement across the population is a 1.1 degree decrease, and the size of the effect increases as baseline auditory spatial ability gets worse. On average, someone with a 5 degree central separation threshold experiences a 0.5 degree (10%) improvement in threshold but someone with a 15 degree central threshold experiences a 3 degree (20%) improvement. The average improvement in performance at the central threshold is 2.2%.

### Modeling

We developed an ideal observer model for our task in order to investigate whether our data are compatible with an optimal combination of auditory and visual cues in this task. Our model (details in Methods) follows Kording et al. [2] in performing inference over whether two cues are the result of the same event, or due to different events (“causal inference”). Cues stemming from the same event are combined according to their relative reliabilities in an optimal manner. This results in a posterior belief about the location of the auditory tone. If this posterior has more mass left of the midline, the ideal observer responds “left”, otherwise “right”.

While the ideal observer performance follows an approximately sigmoidal shape as a function of auditory azimuth as expected, the two model fits corresponding to the matched and central conditions are identical at every angle. The ideal observer’s performance is thus unaffected by the presence of the visual cues and cannot explain the empirically observed behavioral difference between the two conditions.

While a full Bayesian derivation proving that the visual stimuli do not provide a benefit to the ideal observer is given in the Methods, we illustrate a simplified explanation in Figure 3. The subject’s observations imply “initial” subjective beliefs about all four stimulus locations: tone 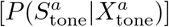, noise 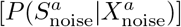, left shape 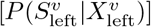, and right shape 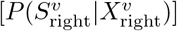. If the brain infers that the auditory and visual stimuli originate from a common source, all four initial beliefs are combined optimally to infer the correct task response (Figure 3). Having learned through task experience that auditory and visual stimuli are always presented symmetrically, the observer can compute a within-modality combined belief, weighting each cue by relative reliability as in Ernst & Banks [1] 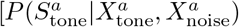 and 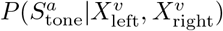 respectively]. Importantly, when combining with the bimodal visual likelihood, the observer must separately consider two possible scenarios: the tone is on the right, or the tone is on the left. Using the visual observation to refine their estimate of the tone location, the observer combines auditory and visual information for each scenario and must base their final decision on a weighted combination of these multisensory beliefs 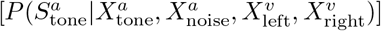. Even weighting the two scenarios equally, there is more evidence in favor of the tone being on the right, the same side as that implied by just the auditory observations. In reality, the weights will depend on the proximity of auditory and visual observations, favoring the visual cue that falls on the same side of the midline as the subject’s belief about the tone and will therefore yield an identical response to the one got by considering just the auditory observations. Equivalently, the side with the greater mass for 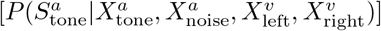 is the same as that for 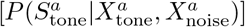. As a result, using the visual stimuli to refine the final posterior does not change the side with more probability mass (Figure 3), and therefore cannot benefit the ideal observer.

**Fig 3.**
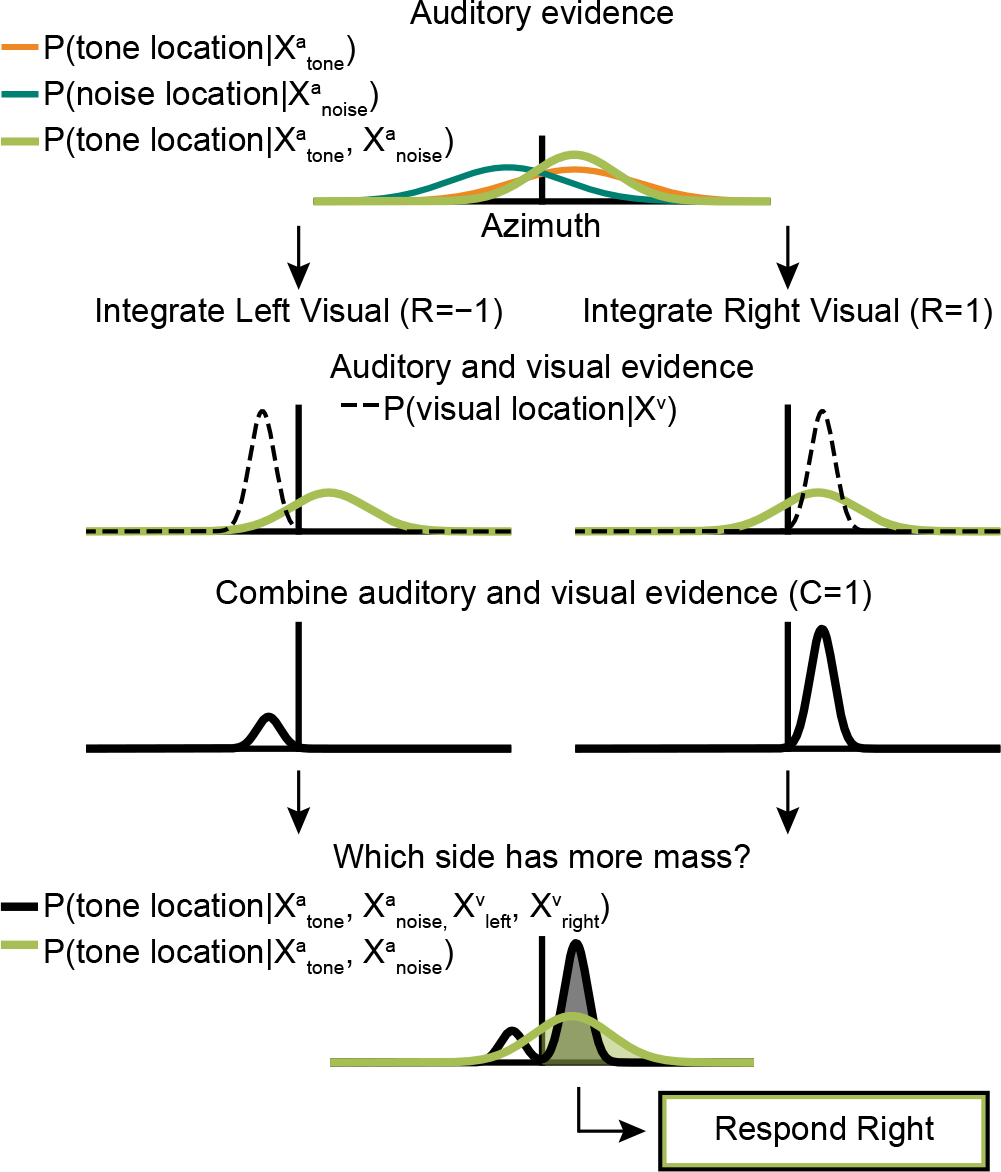
Schematic showing that visual combination cannot provide a benefit to the ideal observer. Listeners use the knowledge that the tone and noise are symmetrically presented to compute a combined auditory likelihood. Then, for each side, they combine this auditory likelihood with a visual likelihood similarly devised from both visual shapes’ likelihoods. Listeners determine which side the tone is on by picking the side of the posterior with more probability mass. Whether they do or do not combine evidence across modalities, the observer responds right.

## Discussion

Here we show that normal hearing listeners improve their performance in an auditory spatial discrimination task when spatially aligned but task-uninformative visual stimuli are present. We further show that these findings cannot be explained by an ideal observer performing the discrimination task.

Even though the shapes presented on any given trial give no indication of which side the tone is on, subjects’ behavioral performance suggests the spatial information they provide can still be combined with auditory cues to reduce errors. Phenomena not captured by the ideal model are needed to explain these results. Assuming that the listener uses the information that the tone and noise are presented symmetrically and bases their decision on the relative positions of the two stimuli, response errors can arise from one of two situations: both auditory stimuli are perceived at the same location (respond at chance), or the relative position of the two auditory stimuli is reversed (response will be incorrect). If the listener only bases their decision on the sign of the position of the tone, errors will occur whenever the tone location estimate crosses the midline. In either scenario, we posit that visual stimuli can act as anchors to attract auditory location. The brain may therefore correct errors in auditory spatial discrimination by refining one or both auditory locations as long as it is able to correctly determine which auditory and visual stimuli to pair. Additional work must be done in order to understand how the brain accomplishes multisensory pairing.

Another interpretation of the visual benefit would be that the visual shapes help direct auditory spatial attention. The time required to shift auditory spatial attention is on the order of 300 ms [15], rendering it unlikely that attention is driving the present results. Visual stimuli preceded the auditory stimuli by 100 ms and the auditory stimuli were only 300 ms long, a duration insufficient for the brain to redirect attention to either of the visual locations, let alone both (splitting auditory attention can have profound behavioral costs [16, 17]).

We find that the ideal Bayesian causal inference model cannot account for the benefit provided by spatially aligned visual stimuli. In particular, we have proven that the visual stimuli in our task can reduce the variance of the auditory estimate but the side on which most of the probability mass lies, and hence the decision, never changes. This means that an ideal observer model is insufficient to explain the benefit we measured behaviorally.

The nature of the suboptimality that allows listeners to benefit from task-uninformative visual stimuli is yet unknown. Different decision strategies for the causal inference model have been tested with traditional audiovisual localization tasks, suggesting that individuals may use a suboptimal probability matching strategy [18]. Such a strategy could also drive the effect we measured. Alternatively, a model of an observer whose low-level auditory coding is biased by visual stimuli (i.e., an early integration model [19]) might also explain the effect. There is some evidence of this type of early integration in the visual cortex during audiovisual localization [12]. Future work exploring such nonideal perceptual mechanisms and their effects on behavior is necessary.

Here we show that listeners use task-uninformative visual stimuli to improve their performance on an auditory spatial discrimination task. This finding demonstrates that the brain can pair auditory and visual stimuli in a more complex environment than typically created in the lab to improve judgments about relative auditory position. The failing of the ideal Bayesian causal inference model to replicate this effect also indicates that these listeners deviate from ideal observers in systematic ways that may lead to insights into the underlying multisensory mechanisms.

## Methods

### Psychophysics

24 Participants (14 female, 10 male) between ages of 19–27 years (mean of 22) gave written informed consent to participate and were paid for time spent in the lab. Each subject had normal or corrected-to-normal vision and normal hearing (thresholds of 20 dB HL or better for octave frequencies between 500 and 8000 Hz). During the experiment subjects were seated in a dark soundproof booth with a viewing distance of 50 cm from a 24 inch monitor with the center of the screen approximately lined up with their nose. Protocol was approved by the University of Rochester Research Subjects Review Board.

#### Stimuli

Two auditory stimuli were generated in the frequency domain with energy from 220 to 4000 Hz and a 1/*f* envelope (−3 dB/octave). One was pink noise (‘noise’) and the other was composed of harmonics of 220 Hz (‘tone’). With the exception of one subject who was run with a frozen noise token, the noise was randomly generated for each trial. Data were similar for the subject with the frozen noise token and therefore not excluded. In order to change the location of each sound, they were convolved with HRTFs (head related transfer functions) from the CIPIC library [20]. Because the experimentally determined HRTFs were only recorded at intervals of 5 degrees in the azimuthal plane, we angles between 0 and 5 degrees were generated from interpolated HRTFs (see expyfun). Adapting methods from ([21]), we generated weights for each of two known HRTFs based on distance from the desired HRTF. Then we took the weighted geometric mean of the known HRTF amplitudes and the weighted arithmetic mean of the angles. After convolution, noise and tone were summed and given a 20 ms raised-cosine ramp at the on and offsets. They were presented at 65 dB SPL at a sampling frequency of 24414 Hz from TDT hardware (Tucker Davis Technologies, Alachua, FL). Auditory stimuli had a duration of 300 ms.

The visual stimuli were regular polygons inscribed in a circle with diameter 1.5 degrees. They were randomly assigned four to eight sides for each trial while ensuring that the two shapes were different. The colors of the shapes were specified according to the HSL scheme, and had constant luminance of 0.6, saturation of 1, and per-trial random hue such that the two shapes in the trial had opposite hue. Each shape was presented during alternating frames at 144 frames per second such that both shapes were visible, even in cases where they would overlap (in a manner similar to [22]).

#### Task

During each trial, the tone and noise were presented symmetrically about zero degrees azimuth with visual onset leading auditory by 100 ms. Subjects were asked to report which side the tone was on by pressing one of two buttons. Before the experiment, subjects were given 10 sample trials and then asked to complete a training session. Their responses to training trials with auditory stimuli at 20 degrees separation were logged until at least 20 trials had been completed and the probability of achieving that number of correct responses by chance (assuming a binomial distribution) was under 5%. If the training criteria was not satisfied after 20 trials subjects were allowed to re-attempt once. Four subjects were dismissed when they did not pass the second attempt.

There were two conditions tested: a matched condition in which the visual and auditory stimuli were spatially aligned, and a central condition in which the visual stimuli were located at the center of the screen (providing no information about the auditory stimuli and therefore serving as a control). Within these conditions we tested five different auditory separations: 1.25, 2.5, 5, 10, and 20 degrees. For each separation there were 80 trials (40 with the target on the right and 40 with the target on the left) for a total of 800 trials. Conditions and separations were randomly interleaved such that the conditions could only lag each other by 2 trials. After subjects got a multiple of 3 trials correct in a row, they were given an encouraging message telling them how many consecutive correct responses they had given. After each set of 40 trials, participants were given a self-timed break.

#### Analysis

We performed maximum likelihood fits to the percent correct of the responses at log transformed auditory separations. First we estimated the lapse rate and slope of each subject by doing a preliminary sigmoidal fit on the pooled responses to both conditions. Then using these estimates of lapse rate and slope, we fit responses for both conditions, central (control) and matched, only letting midpoint vary. The lapse rate and slope should be independent of the visual condition. Thresholds were approximated as the separation level at which the fit crossed 75% correct. Using *p* < 0.05 as the criteria for significance, we compared the matched and central percent correct measures with paired *t*-tests. Because thresholds were not normally distributed across subjects, changes thereof were assessed with a sign test.

### Modeling

We model the subject responses from a normative perspective by using an ideal observer model. The subjects are assumed to have learned a generative model of the inputs and base their decision on the inferred tone side. The structure of the model is generally summarized in Figure 4.

**Fig 4.**
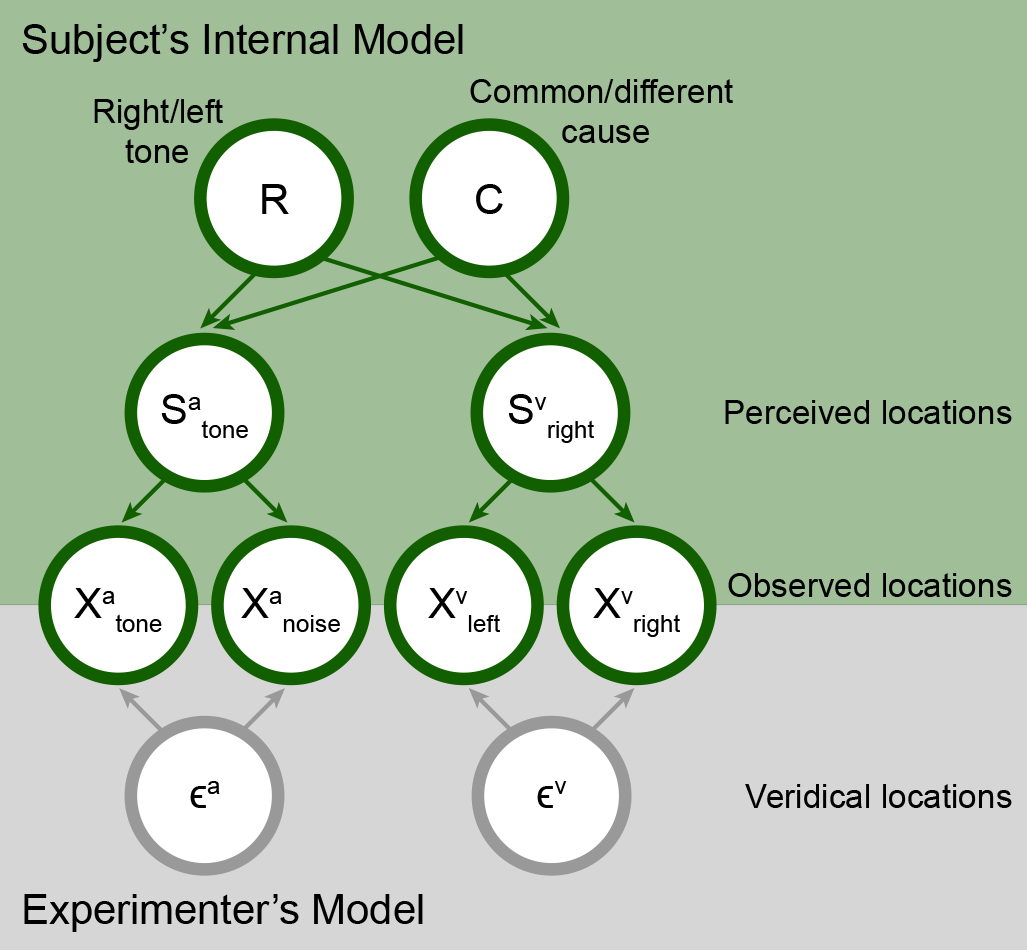
Graphical depiction of our model schematic. Our full model contains two generative models. The first one is the experimenter’s model which maps the true task variables to the sensory observations made by the subject. The second is the subject’s internal model of the sensory observations which is used the subject’s perception (Inference in the generative model).

#### Model definition for a single trial

For each trial, we denote the true auditory tone location (signed) as 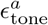 and true visual cue eccentricity (always positive) as 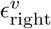 (this is sufficient to define all the inputs since the true noise location and true left visual cue locations are the negatives of the aforementioned values). In this notation, 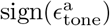 denotes the correct response for that trial. Using the notation 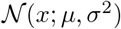 to denote the probability density function of a normal random variable with mean *μ* and variance *σ*^2^, the observed tone location 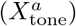, noise location 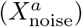, left visual cue location 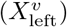 and right visual cue location 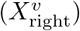 for the trial are randomly drawn with probability:

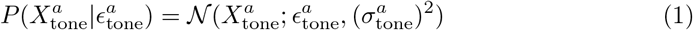

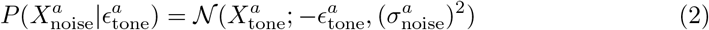

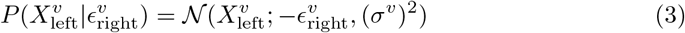

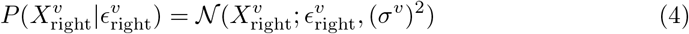

where 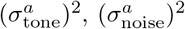, (*σ*^*v*^)^2^ are the uncertainties associated with the observed tone, noise and visual cue locations respectively.

It is important to note that the subject does not have access to the true variables 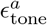 and 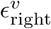 and must make their decision from the observed variables.

We model subject perception as inference in a hierarchical generative model of the sensory inputs (shown in the figure). Let 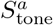 and 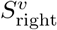 be the perceived tone and right visual cue location whose likelihood are given as follows

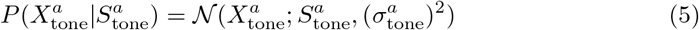

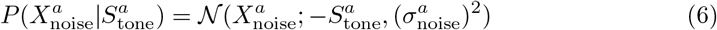

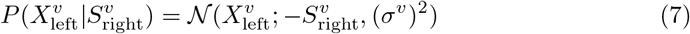

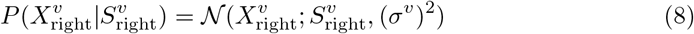

Eq. 5 to 8 assume that the subjects can account for their uncertainty accurately based on prior sensory experience. We assume that the subject has learned that the auditory and visual stimuli are symmetric about zero degrees azimuth, which allows them to collapse 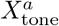 (or 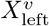) and 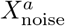 (or 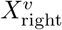) into unimodal estimates. The priors over 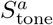 and 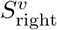 can be conditioned on whether the subject perceived the tone to be from left or right (denoted as R=−1 or R=1 respectively) and if they perceived the auditory and visual cues to be from the same cause or not (denoted by C=1 or C=0 respectively). Assuming a flat prior over location for 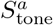 and 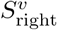(the results still hold for symmetric proper priors), this can be written as

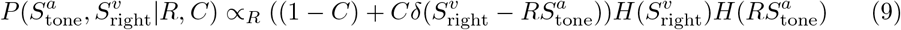

where ∝_*R*_ indicates that the proportionality context is independent of R. *H*(*x*) denotes the Heaviside function.

Having inferred R, we note that the ideal observer makes their choice (Ch) by choosing the side with the higher posterior mass, i.e.

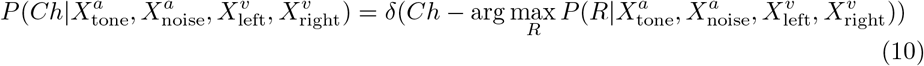

#### Calculating the posterior

Before comparing the probability mass on either side, we must evaluate the posterior over R. In order to do so, we marginalize over the cause variable C

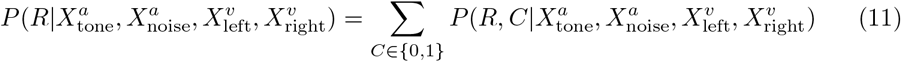

We can evaluate the term inside the sum by first using Bayes rule and then simplifying under the assumption that the priors over R and C are assumed to be independent, i.e. *P* (*R, C*) = *P* (*R*)*P* (*C*)

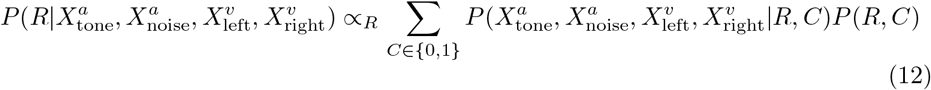

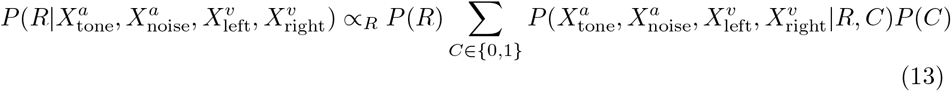

By assuming equal priors for the left and right side, i.e. *P* (*R*) = 0.5(Ideal observer has no response bias.

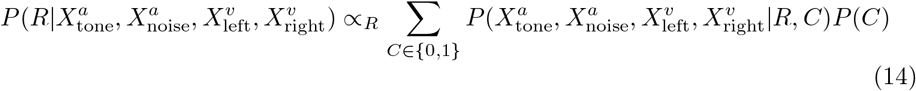

We can then expand the expression of the side with the higher posterior mass by considering both values of the cause variable C, which using Eq. 14 can be rewritten as

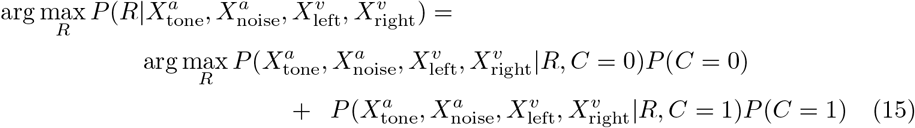

In general, the likelihood can be evaluated by averaging over all possible auditory and visual cue locations

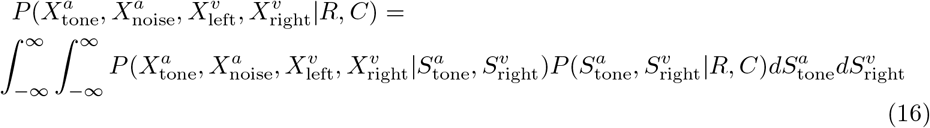

Using the independence relations implied by the generative model, we can simplify the previous equation to get

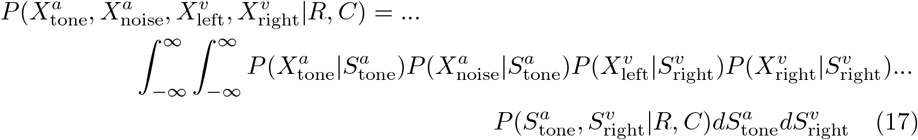

Substituting expressions for the likelihoods of each cue (Eq. 5-8) and the prior (Eq. 9), we can evaluate Eq. 17 by repeated multiplication of normal probability density functions to get expressions for both *C*=0 and *C*=1

#### No audiovisual combination (C=0)

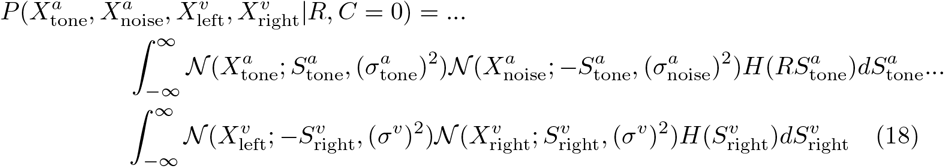

Multiplying the gaussian likelihoods in Eq. 18, we can pull the terms independent of R into a proportionality constant to get

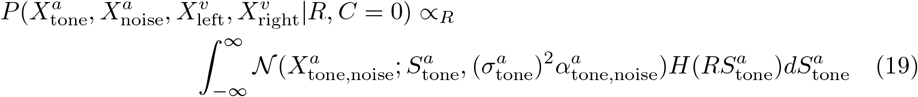

where

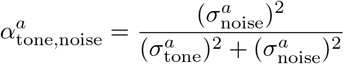

is the weight given to the tone location while combining with the noise location.

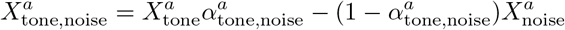

is the combined estimate of the auditory tone location by weighting the tone and noise observation by their inverse variances. The integral in Eq. 19 is the area of the combined gaussian likelihood for the tone and noise on either the positive or negative side of 0 depending on R.

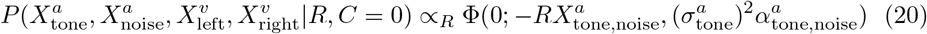

where Φ(*x*; *μ, σ*^2^) denotes the cumulative density function evaluated at x for a normal random variable with mean *μ* and variance *σ*^2^.

#### Audiovisual cue combination (C=1)

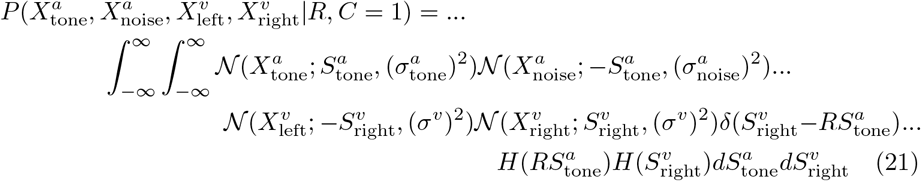

We can integrate over 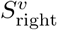 by evaluating all functions of 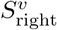 at 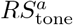 because of 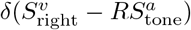

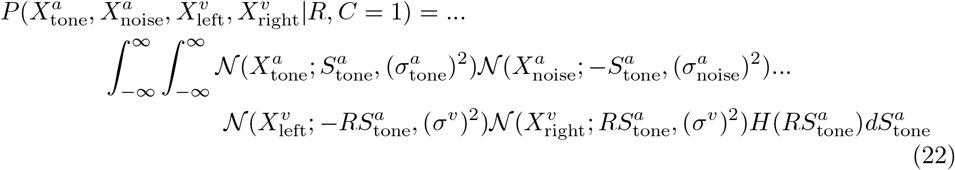

Multiplying the gaussian likelihoods in Eq. 22, we can pull the terms independent of R into a proportionality constant to get

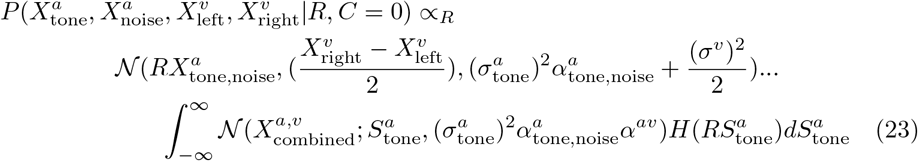

where

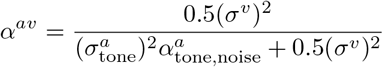

is the weight given to auditory location while combining with the visual location and

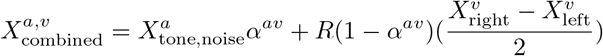

is the weighted combination of the visual and auditory cues. The integral in Eq. 23 evaluates to

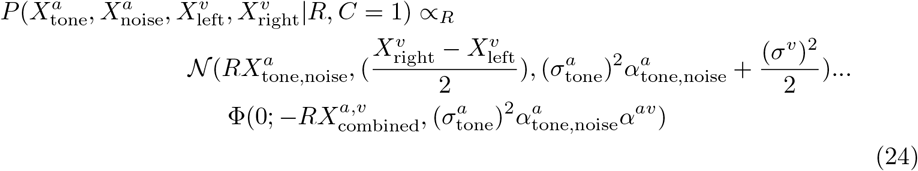

Using the fact that Φ(0; *μ, σ*^2^) is a decreasing function of *μ*, the maximum of Eq. 20 simplifies to

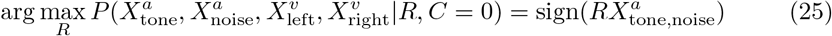

We note that 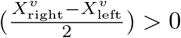 (by definition). Using that fact 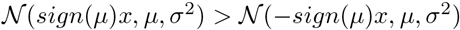 in addition to the decreasing nature of Φ(0; *μ, σ*^2^), the maximum of Eq. 24 simplifies to

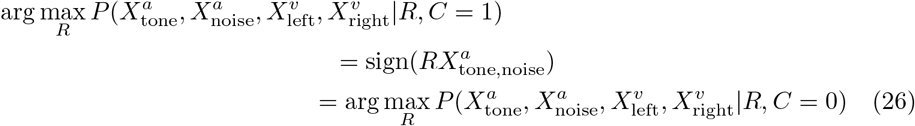

The positive weighted combination of two function is maximized at the point of maximization of the individual functions if the individual point of maximizations are equal. Using this result, we can substitute Eq. 21 into Eq. 15 to get

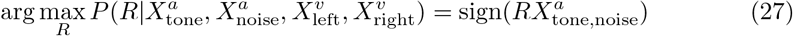

Importantly, the side with the higher posterior mass is independent of cause C.

#### Generating a psychometric curve

To evaluate the probability the subject will choose right at each auditory azimuth (psychometric curve), we need

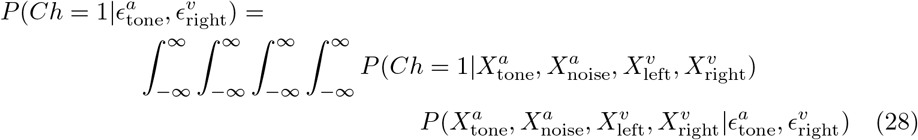

Using the independence relations implied by the generative model, we can simplify the previous equation to get

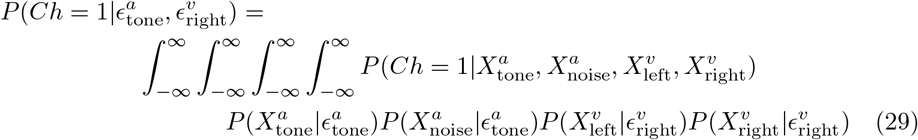

Substituting Eq. 1-4, 27 in Eq. 29 and simplifying, we get

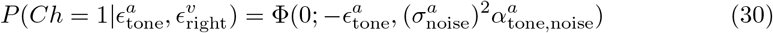

Assuming a subject lapses with a probability λ and responds randomly with equal probability, we get the model predicted psychometric curve as

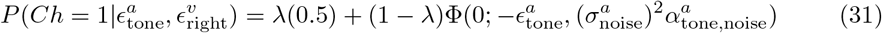

It is important to note that the ideal observer response is **not affected** by the observations of the visual cue location.

### Model Fitting

The model described in the previous section has two free parameters to model the subject responses:

- Effective auditory uncertainty 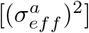
- Lapse rate (λ)

Because the ideal observer is not affected by visual cues, we do not fit a parameter for visual uncertainty. We compute the posterior over these parameters (denoted as *θ*) from subject responses. Let *n*_condition_ denote the number of stimulus conditions in the experiment and *n*_trial_ denote the number of trials for each condition. We denote the true auditory eccentricity and true right visual cue eccentricity for condition *i* as 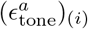 and 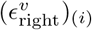 respectively. The experimental subject responses for these conditions are denoted by (*r*)_(*i*)_ which is modeled as a binomial random variable

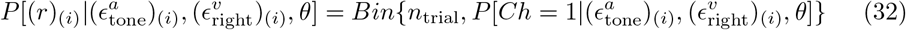

where *Bin*(*n, p*) denotes the binomial probability density function with parameters *n* and *p*. The probability parameter in Eq. (27) is obtained from Eq. 31 for the parameter values.

Given these data points from the experiment, we are interested in calculating the probability of the parameter value given this data, i.e. 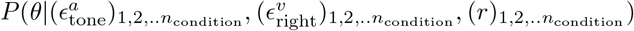. Using Bayes rule,

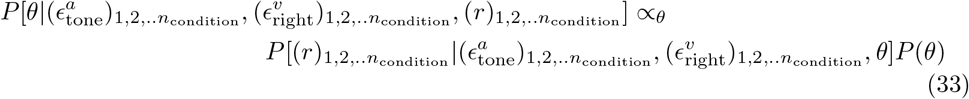

We have assumed that the probability of the subject’s parameters are independent of the cue location. Assuming all conditions are independent given the parameter value (which is assumed to have a flat prior), we simplify Eq. 33 to get

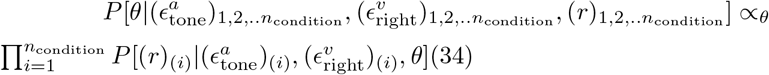

where the term inside the product is given in Eq. 32

We can find the parameters that best fit the data (denoted as *θ*\*) by finding the maximum a posteriori (MAP) solution for Eq. (Also the maximum likelihood since the prior is flat). This is often implemented as minimizing the negative log posterior (since log is monotonic)

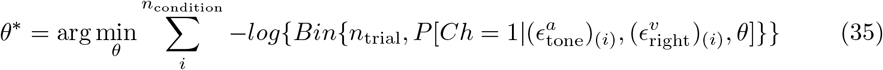

We optimized Eq. 35 using Bayesian adaptive direct search (BADS) [23]. BADS alternates between a series of fast, local Bayesian optimization steps and a systematic, slower exploration of a mesh grid.

## Data and Code

Data and code are available from https://github.com/Haefnerlab

## Acknowledgments

The authors wish to thank Veronica Valencerina for assistance with data collection. This work was supported by NIH grant R00 DC014288 awarded to RKM.

